# Maternal effects regulate population genetic structure by altering intraspecific competitive relationships

**DOI:** 10.1101/2025.09.28.678971

**Authors:** Jia Na, Cuijuan Niu

## Abstract

1. The transgenerational plasticity of maternal effects on offspring phenotypes has been well recognized, however, weather maternal effect can drive dynamic changes in population genetic structure by modifying the competitive dynamics among offspring is still unclear.
2. After evaluation of the reliability of COI metabarcoding for intraspecific clone quantification using a monogonont rotifer *Brachionus calyciflorus* as the model animal, a high-frequency dynamic monitoring experimental system was set to measure maternal effect on temporal dynamics in gene frequencies within mixed populations containing morphologically identical clones.
3. Results showed that maternal effects significantly enhanced the competitive fitness of the target offspring clone, leading to a rapid increase in its frequency inside the population. However, this advantage last for a few generations, and gradually diminished or even reversed as the maternal effects attenuated across generations.
4. Temperature dynamically affected the duration of maternal effect: higher temperature shortened the duration of maternal effects on population genetic dynamics, while lower temperature prolonged it.
5. By population dynamic monitoring, this study revealed the transient nature of maternal effect as a non-genetic buffering mechanism, providing an integrative theoretical framework for understanding the dynamic integration of short-term plasticity and long-term selection pressure in adaptive evolution of intraspecific clones.

## 1. Introduction

As a transgenerational expression of phenotypic plasticity, maternal effects refer to the phenomenon whereby maternal phenotype and environmental experience modulate offspring fitness through non-genetic pathways(Mousseau & Fox, 1998, Wolf & Wade, 2009). In natural ecosystems, multidimensional factors can induce maternal effects that significantly alter offspring phenotype, fecundity, and survival(Richards, 2006, Liebers *et al*., 2014, Uller *et al*., 2015, Feiner *et al*., 2022). Furthermore, maternal effect can persist to grand-offspring and even span multiple generations(Yin *et al*., 2015, Sentis *et al*., 2018, Yin *et al*., 2019, Colinas *et al*., 2023).

Modern population ecology emphasizes that individual variation is central to elucidating population dynamics (Forsman & Wennersten, 2016, Delgado *et al*., 2018). Offspring trait differentiation arising from maternal effects (including survival rate, growth rate, and fecundity) has been demonstrated to regulate population numerical dynamics (Benton *et al*., 2008, Love *et al*., 2012, Moore *et al*., 2019, Xu & Niu, 2021). Regarding population genetic structure, maternal effects may exert equally pivotal functions. Firstly, by conveying environmental cues via epigenetic modifications, maternal effects may buffer the impact of natural selection on population allele frequencies (Yamamichi & Hoso, 2017). Secondly, this plasticity may facilitate offspring in transcending parental phenotypic limitations and broadening ecological niche width (Schneider & Meyer, 2017)and consequently shaping population genetic diversity (Moore *et al*., 2019).

Notably, current researches focus on maternal effects influencing numerical dynamics in monoclonal populations, or on maternal effects regulating entire multiclonal populations, yet fail to clarify a possible role of maternal effects in altering population genetic structure by modulating competitive results among clones. A key scientific question remains unresolved: how maternal effects could modify fitness of particular clonal lineages, thereby inducing intraspecific competition shifts, ultimately driving restructuring of population genetic structure (e.g., clonal frequencies). This core scientific question links the micro-scale mechanisms of maternal effects (monoclonal fitness) with macro-scale evolutionary outcomes (genetic restructuring), holding significant theoretical importance for revealing maternal effects’ role in regulating population dynamics and genetic structure.

The monogonont rotifer *Brachionus calyciflorus* provides an ideal model for addressing this question: parthenogenetic reproduction enables construction of homogeneous clonal lineages that inherently avoid interference from sexual recombination, brief generation times allow accelerated population establishment, and high sensitivity to environmental variations with multidimensional response capabilities (Gilbert, 2004, Gilbert & Schröder, 2004). With intensifying global warming and extreme weather events, temperature as the most direct and drastically changing environmental factor, has received substantial focus for its biological consequences. Number of studies have demonstrated the effects of maternal temperature environment on offspring fitness across diverse organisms. However, how temperature-induced maternal effect influences population genetic structure remains unclear, in spite of its importance for predicting adaptive evolutionary pathways.

A central methodological challenge is the dynamic quantification of intraspecific clonal frequencies. Traditional morphological identification struggles to achieve clonal-level identify due to high phenotypic similarity among clones, while microsatellite genotyping can provide genetic marker information it suffers from limited detection throughput (Lemmen *et al*., 2023). Although DNA metabarcoding is widely used in community analysis(Hebert *et al*., 2003, Zaiko *et al*., 2015, Serrana *et al*., 2022), its reliability in quantifying clonal proportions at sub-species level remains doubtful because of PCR amplification biases and sequence homology among clones with minimal genetic divergence that may introduce substantial quantification errors (Elbrecht & Leese 2015; Clarke *et al*. 2017; Serrana *et al*. 2022). Therefore, systematically evaluating the reliability boundaries of DNA metabarcoding for clonal proportion quantification becomes the primary task in related experimental design.

In this study, we collected *B. calyciflorus* from natural habitats and established multiple monoclonal populations through laboratory acclimation.We then sequenced their COI barcodes and selected two maximally divergent clones (S and A; COI p-distance = 0.53) to conduct the following experiments: (1) Artificial construction of populations with predefined clonal ratios to evaluate COI metabarcoding accuracy in resolving clonal proportions; (2) Maternal temperature effects (high/low temperature treatment) were applied to clone S, and monitoring changes in competitive interactions with the control clone A. We quantified the impact of maternal effects on population genetic structure by analyzing numerical dynamics of mixed clonal populations and temporal changes in clonal proportions. We hypothesized that maternal environmental will alter fitness of offspring, reshape competitive relationships between the clones, and thereby drive adaptive changes in clonal frequency within populations over time. Our research challenges the traditional paradigm that “maternal effects act solely as ecological modifiers,” pioneering the recognition of their functions as evolutionary drivers of population genetic structure dynamics. It will establish a theoretical framework for forecasting trajectories of adaptive evolution in the area of global changes.

## 2. Materials and Methods

### 2.1. Cultivation and identification of organisms

Clonal lineages A and S of the rotifer *B. calyciflorus* utilized in this study, were both collected from Lake Aohai of Olympic Forest Park, Chaoyang District, Beijing (40.018°N, 116.404°E), long-termly cultivated in the laboratory stable environment with resting egg production. At experiment initiation, one resting egg was hatched to establish monoclonal populations. Rotifers of different clones were cultured in COMBO medium (Kilham *et al*., 1998) at 20°C with 16:8h light: dark photoperiod in incubators. During acclimation, medium containing *Chlorella pyrenoidosa* was refreshed daily with random removal of rotifers to maintain densities at 0.1 inds ml^-1^(rotifers) and 1×10^6^ cells ml^-1^ (algae), continuing >30 days before formal experiments. The algal strain *C. pyrenoidosa* (FACHB-5) was purchased from the Freshwater Algae Culture Collection at the Institute of Hydrobiology,Chinese Academy of Sciences. Algae were grown in BG-11 medium under aerated conditions in incubators, then centrifuged (4°C, 7000 rpm, 7 min) at peak vitality. Cell pellets were resuspended in COMBO medium to generate concentrated algal stocks, then were enumerated microscopically and refrigerated at 4°C until required. Working solutions were prepared by dilution to specified concentrations prior to application.

The mtCOI region and nuITS1 regions were sequenced and assembled using bidirectional Sanger sequencing for each clone. Amplification of COI fragments utilized primer pair LCOmodBc/HCOmodBc, ITS amplification employed primers III/VIII, with both primer sequences and PCR cycling conditions matching those described by (Papakostas *et al*., 2016). No ITS sequence variation was detected between clones, confirming their classification as *B. calyciflorus s*.*s*. in the cryptic species complex(Papakostas *et al*., 2016). Across the 600-bp overlapping COI region, we detected 32 single-nucleotide polymorphisms (SNPs) with a calculated p-distance of 5.30%. All consensus sequences were submitted to the NCBI GenBank database. GenBank accession numbers for Clone S are PP968173(COI) and PV574018 (ITS), Clone A are PP968402 (COI) and PV577650 (ITS).

### 2.2. Confirmation high-Throughput sequencing accuracy in quantifying rotifer clonal proportions

For assessing high-throughput sequencing (HTS) precision in quantifying clonal structure, mixed-clone samples with predetermined proportions were prepared through microscopic counting. Experimental treatments (500 individuals per replicate) covered seven target S-to-A ratios: 20:1, 10:1, 5:1, 1:1, 1:5, 1:10, and 1:20. Stereomicroscope-guided sorting yielded exact compositions: S: A = 476:24, 454:45, 417:84, 250:250, 84:417, 45:454, and 24:476 across treatments. Resulting measured S-to-A ratios were 19.83, 10.09, 4.96, 1.00, 0.20, 0.10, and 0.05 respectively. Following sorting, samples underwent filtration and DNA extraction, with subsequent PCR, high-throughput sequencing, and bioinformatic analysis.

### 2.3. Impact of clonal maternal effects on mixed-population genetic structure

For examining maternal effects in clonal competition dynamics, mix-cultures were initiated using clones A and S of the rotifer *B. calyciflorus*. Maternal effect manipulations (details in 2.3.1) were applied to clone S prior to mixing with control clone A(untreated). Temporal sampling of mixed clone populations enabled DNA extraction and sequencing, with clonal ratios determined through HTS analysis at each interval.

#### 2.3.1 Impact of maternal high-temperature exposure on offspring competitive dynamics among rotifer clones

F0 individuals (neonates <4h) from acclimated clone S were distributed to either thermal stress (28°C, named H) or control (20°C, named C) environments and reared to sexual maturity. The first amictic female offspring from F0 became F1 generation; offspring from C- and H-reared mothers were randomly assigned to matching or mismatching temperatures, forming four treatments: CC, HC, HH, CH (first letter: F0 temperature; second: F1/rearing temperature). Clone A (neonates <4h) were reared as F0 at control temperature (20°C), with their first amictic female offspring (F1) subsequently allocated to all four treatment groups. Each replicate was initiated with 5 maternal-effect treated clone S amictic females and 5 untreated clone A amictic females, with daily renewal of food-enriched medium. The mixed-clone populations were cultured continuously for 20 days, with daily counts, differentiation of female types and recording of resting egg production and then collection of resting eggs. Fifteen replicates per treatment were established initially; triplicates were filtrate at 4-day intervals (resting eggs removed prior to filtration), preserved at -20°C for DNA analysis (Detail can be found in 0), with demographic data collected solely from three fixed replicates. All culture conditions except temperature were maintained identical to the acclimation protocols described in Section2.1 throughout the experiment. Fig. 1 shows a detailed schematic of the experimental procedures.

**Fig. 1.**
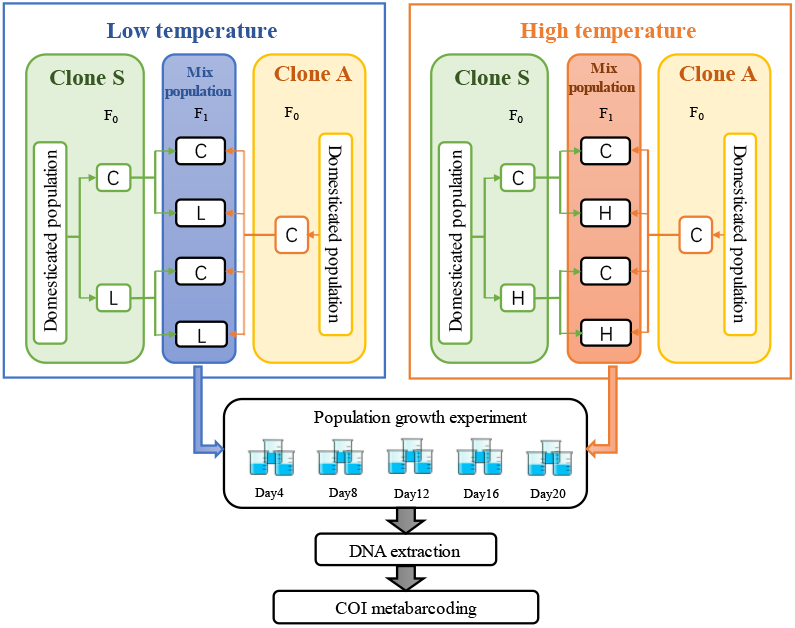
Experimental design for maternal effects on genetic structure in mixed clonal populations. The two-letter notation specifies maternal (F0) environment (first letter) and offspring environment (second letter). C denotes control (20°C), H denotes high temperature (28°C), and L denotes low temperature (12°C).

#### 2.3.2 Impact of maternal low-temperature exposure on offspring competitive dynamics among rotifer clones

Experimental design mirrored Section 0, with thermal exposure replaced by cold exposure (12°C, named L), generating treatment groups CC, LC, LL, CL.

### 2.4. Sample preprocessing and DNA extraction

Samples filtrated through 1.2 μm MF-Millipore™ MCE membrane(Millipore, Tianjin). Contamination control involved soaking filtration equipment and forceps in 10% 84 disinfectant (30 min), with subsequent washing in tap water and distilled water. Prior to filtration, benches were sanitized with 10% 84 disinfectant and ethanol, with equipment changed between samples to avoid cross-contamination. Samples were decanted into filtration assemblies with gentle swirling, followed by COMBO medium washes to ensure complete sample deposition on filters. Process controls (COMBO medium blanks) were filtered concurrently with experimental samples.

Membranes were transferred to 2 mL sterile tubes after filtrating, sealed and cryopreserved at - 20°C. Prior to DNA extraction, membranes were minced in-tube using disinfected (84 disinfectant -soaked) and rinsed scissors. DNA was extracted using an optimized DNeasy Blood and Tissue Kit (QIAGEN, Germany) procedure(Zhang *et al*., 2020). DNA concentration was measured with a NanoDrop One system (Thermo Fisher Scientific, USA). PCR templates were prepared as 5× dilutions to minimize amplification inhibition (Zhang *et al*., 2022).

### 2.5. Primer and PCR amplification

In zooplankton metabarcoding studies using high-throughput sequencing, mitochondrial cytochrome c oxidase subunit I (COI) is the most extensively employed target fragment and exhibits the highest species discriminatory power. The mlCOIintF/jgHCO2198 primer pair shows superior amplification performance and taxonomic resolution for invertebrates(Elbrecht & Leese, 2017, Leray *et al*., 2013). Across this 313-bp amplicon, interconal variation between clone S and clone A comprised 18 SNPs with 5.75% sequence divergence (Appendix). Based on published amplification efficacy and observed sequence divergence, primers mlCOIintF (5’-GGWACWGGWTGAACWGTWTAYCCYCC -3’) and jgHCO2198 (5’- TAIACYTCIGGRTGICCRAARAAYCA -3’) were chosen for 313-bp COI amplification(Leray *et al*., 2013).To distinguish PCR reactions, 7-bp unique barcodes were appended to the 5’ termini of each primer (Coissac, 2012, Zhang *et al*., 2020). Barcode designs are detailed in the Appendix.

The 25-μL PCR reaction mixture comprised: 2.5 μL DNA template, 1 μL (0.2 μM) of each forward and reverse primer, 12.5 μL 1× Premix Ex Taq (Takara, Kusatsu, Japan), with ddH^2^O added to a final volume of 25 μL. PCR amplification was performed under the following conditions: 94°C for 2 min; 30 cycles of 94°C for 1 min, 55°C for 1 min, and 72°C for 1 min; with a terminal extension at 72°C for 10 min and cooling to 4°C for storage. Each DNA sample was amplified in triplicate, accompanied by 5 filtration blanks, 1 extraction blank, 5 negative controls, and 2 positive controls per PCR batch. Following quality assessment via 1% agarose gel electrophoresis, amplicons were normalized, pooled, and transferred to Novogene Corporation (Beijing, China) for sequencing. Library sequencing employed 250-bp paired-end reads (PE250) in PCR-free mode on an Illumina NovaSeq 6000 system (Novogene Corporation, Tianjin, China).

### 2.6. Bioinformatics analysis

Raw data were processed using the OBITools pipeline(Boyer *et al*., 2016). Primary sequence alignment paired forward and reverse reads via the illuminapairedend module. Reads exhibiting Phred scores below 40 were eliminated through obigrep filtering. Sequence selection via ngsfilter required exact tag matches (0 mismatches) and permitted ≤2 primer mismatches. dentical sequences were then clustered into unique sequence variants using obiuniq. Low-abundance sequences (<10 total reads) and short fragments (<150 bp) were excluded through additional obigrep filtering. Artifactual sequences from PCR/sequencing errors were detected and eliminated with obiclean (abundance ratio threshold=0.5; sequence difference=1)(Schnell *et al*., 2015).

Processed sequences were aligned against a custom reference database (containing COI sequences from clone S and clone A obtained in Section 2.1) using BLASTN. Annotation outputs within 97-100% similarity thresholds were comparatively assessed for the validation experiment (Section2.2). In the maternal effect experiment (Section 2.3), only annotations with ≥98% sequence similarity were retained. Best-match assignments were derived by processing BLASTN outputs through jcvi.formats.blast (Tang *et al*., 2024). Data were normalized against contamination using the formula: Final read count = Raw sample count - Maximum count in concurrent negative controls, thereby mitigating cross-contamination and tag-jumping effects (De Barba *et al*., 2014). The complete bioinformatic processing pipeline code can be found at https://pythonhosted.org/OBITools/welcome.html, and detailed statistical data on numbers of sequence reads for each library at each stage are presented in the Appendix.

### 2.7. Statistical analysis

All data analyses and visualization were implemented in R (version 4.4.2). To validate sequencing quantification accuracy, the mixed-clone confirmation experiment employed linear fitting models (lm) to test statistical concordance between clone S/A sequencing reads ratios and actual composition of biological samples.

To assess maternal effects on mixed-clone population genetic structure, generalized estimating equations (GEE) modeled variations in population density, proportion of mictic females, and resting eggs production. Temporal dynamics of the S/A clone ratio were analyzed via two-way factorial analysis using either parametric ANOVA (for normally distributed, homoscedastic data) or Scheirer-Ray-Hare’s rank-based method (for non-normal/heteroscedastic distributions). These models tested the effects of maternal and offspring environments on population genetic structure. Tukey’s Honest Significant Difference (HSD) test conducted post hoc pairwise comparisons between treatment groups.

## 3. Result

### 3.1. Confirmation high-throughput sequencing accuracy in quantifying rotifer clonal proportions

In the validation experiment, sequence proportions of clones S and A were obtained by aligning high-throughput sequencing (HTS) data against our custom database at 97-100% sequence similarity thresholds. Linear regression between these proportions and actual clone size numerical ratios demonstrated R^2^ > 0.970 and P < 0.001 across all similarity thresholds. This indicates excellent goodness-of-fit and a highly significant linear relationship (Fig. 2).

**Fig. 2.**
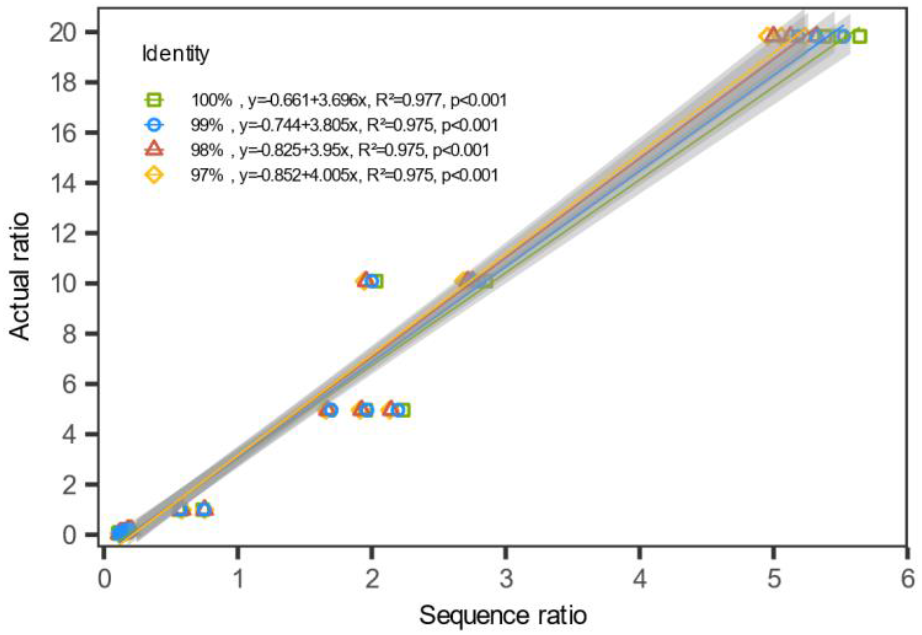
Results of linear regression (lm) between actual S/A clone ratios and COI metabarcoding sequence reads proportions from the validation assay. Colored regression lines in the figure denote sequence similarity thresholds during database alignment. Shaded bands surrounding lines indicate 95% confidence intervals for the regression models.

### 3.2. Impact of clonal maternal high-temperature exposure on mixed-clone population dynamics and genetic structure

Statistical analysis showed maternal environmental temperature did not affect population density, mictic ratio, or daily resting eggs production (P > 0.05, Table 1). Although offspring environmental temperature also did not affect population density (P > 0.05,Table 1), it significantly influenced both mictic ratio (P = 0.012,Table 1) and daily resting egg production (P = 0.003,Table 1). No significant interactions between maternal and offspring environmental temperatures were detected for any of the three examined variables (Table 1). Among the four treatment groups comprising heat-exposed maternal S-clones and non-exposed A-clones, mixed-clone populations with identical offspring temperatures (CC vs. HC; HH vs. CH) exhibited similar temporal variation trend in population density, mictic ratio, and daily resting egg production (Fig. 3). Population density curves showed CH and HH exhibiting elevated population densities relative to CC/HC treatments until day 11, after which CC/HC groups surpassed them (Fig. 3a). Both mictic ratio and daily resting egg production reached peak values earlier in CH/HH groups than in CC/HC treatments, though with lower peak values (Fig. 3b, c), but the HH achieved higher peak values than CH (Fig. 3c).

**Table 1.** Results of the GEE two-way ANOVA on quantitative dynamic characteristics (population density, mictic ratio, daily resting eggs production) of mixed clonal populations in the high-temperature maternal effects experiment.

|  | a) Population density |  |  |  | b) Mictic ratio |  |  |  | c) Number of resting eggs |  |  |  |
| --- | --- | --- | --- | --- | --- | --- | --- | --- | --- | --- | --- | --- |
|  | Estimate | Std Error | Z value | P value | Estimate | Std Error | Z value | P value | Estimate | Std Error | Z value | P value |
| Maternal environment(A) | -4.949 | 3.820 | 1.676 | 0.195 | -0.007 | 0.041 | 0.031 | 0.861 | 11.500 | 21.100 | 0.298 | 0.585 |
| Offspring environment(B) | -0.149 | 3.610 | 0.002 | 0.967 | -0.091 | 0.036 | 6.277 | 0.012 | -45.700 | 15.200 | 9.058 | 0.003 |
| A×B | 3.465 | 4.660 | 0.554 | 0.457 | 0.025 | 0.052 | 0.221 | 0.638 | 2.100 | 23.500 | 0.008 | 0.929 |

**Fig. 3.**
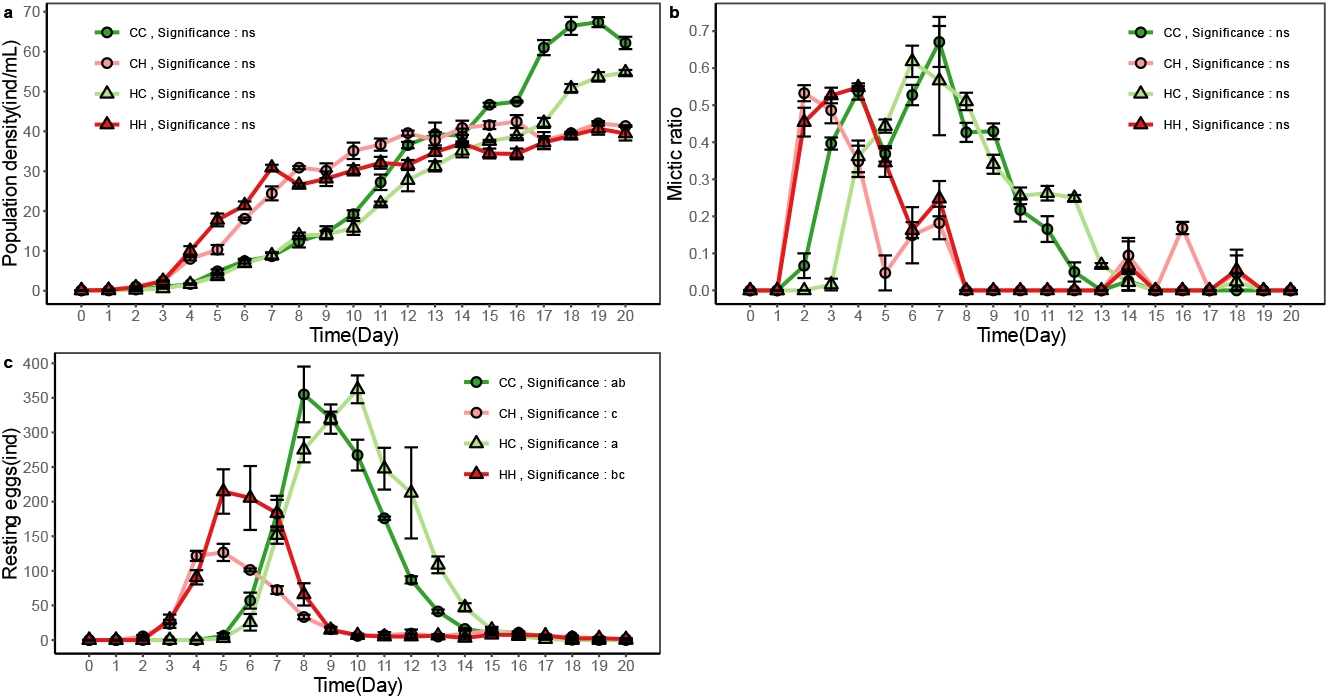
Population dynamics of mixed-clone populations under maternal thermal effects in high-temperature experiments. a: Temporal changes in population density; b: Mictic ratio (= mictic females / (mictic females + amictic females)); c: Daily resting eggs production. In the two-letter codes, the first capital letter indicates the maternal environment, while the second indicates the offspring environment. C represents control temperature (20°C), and H represents high temperature (28°C). Curves labeled with different lowercase letters differ significantly (P < 0.05) as determined by GEE modeling. Data presented as mean ± SE.

Statistical analysis indicated that maternal environmental temperature significantly influenced clone ratios only at day 16 (P = 0.03,Table 2). Offspring environmental temperature exerted significant impacts on clone ratios at days 4 and 8 (P < 0.001,Table 2). However, significant interactions between maternal and offspring environmental temperature were detected on days 4 and 12 (P < 0.001,Table 2). Post hoc Tukey HSD tests showed that clone ratios at day 4 in CC group were significantly higher than HC (P < 0.05), while HH group were markedly higher than HC (P < 0.05,Fig. 4). At day 8, no significant differences were found between CC and HC or between HH and HC (P > 0.05, Fig. 4). However, combined CC and HC ratios were significantly lower than those of HH and HC proportions (P < 0.05,Fig. 4). At day 12, clone ratios of CH group were clearly higher than the other three groups (P < 0.05,Fig. 4). At day 16, no significant differences in clone ratios were observed among all groups (P > 0.05,Fig. 4). Patterns at day 20 mirrored those at day 8. No significant differences were detected between CC and HC groups except at day 4 (P < 0.05,Fig. 4).

**Table 2.** Results of two-way ANOVA on the proportional representation of S-clone and A-clone COI metabarcoding sequences at five time points in the high-temperature maternal effects experiment. F-value represents the parametric two-way ANOVA statistic, whereas H indicates the non-parametric Scheirer-Ray-Hare.

|  | a) Ratio of clone S to A in day 4 |  |  |  | b) Ratio of clone S to A in day 8 |  |  |  | c) Ratio of clone S to A in day 12 |  |  |  |
| --- | --- | --- | --- | --- | --- | --- | --- | --- | --- | --- | --- | --- |
|  | df | Sum sq | F value | P value | df | Sum Sq | H | P value | df | Sum Sq | H | P value |
| Maternal environment(A) | 1 | 1.243 | 0.523 | 0.475 | 1 | 13.444 | 0.121 | 0.728 | 1 | 324.000 | 2.919 | 0.088 |
| Offspring environment(B) | 1 | 47.055 | 19.801 | <0.001 | 1 | 2916.000 | 26.270 | <0.001 | 1 | 205.444 | 1.851 | 0.174 |
| A×B | 1 | 129.876 | 54.653 | <0.001 | 1 | 113.778 | 1.025 | 0.311 | 1 | 1600.000 | 14.414 | <0.001 |
| Residuals | 32 | 76.044 |  |  | 32 | 841.778 |  |  | 32 | 1755.556 |  |  |
|  | d) Ratio of clone S to A in day 16 |  |  |  | e) Ratio of clone S to A in day 20 |  |  |  |  |  |  |  |
|  | df | Sum Sq | H | P value | df | Sum Sq | H | P value |  |  |  |  |
| Maternal environment(A) | 1 | 1002.778 | 9.034 | 0.003 | 1 | 32.111 | 0.289 | 0.591 |  |  |  |  |
| Offspring environment(B) | 1 | 93.444 | 0.842 | 0.359 | 1 | 2916.000 | 85.889 | 26.270 |  |  |  |  |
| A×B | 1 | 5.444 | 0.049 | 0.825 | 1 | 128.444 | 20.193 | 1.157 |  |  |  |  |
| Residuals | 32 | 2783.333 |  |  | 32 | 808.444 | 3.225 |  |  |  |  |  |

**Fig. 4.**
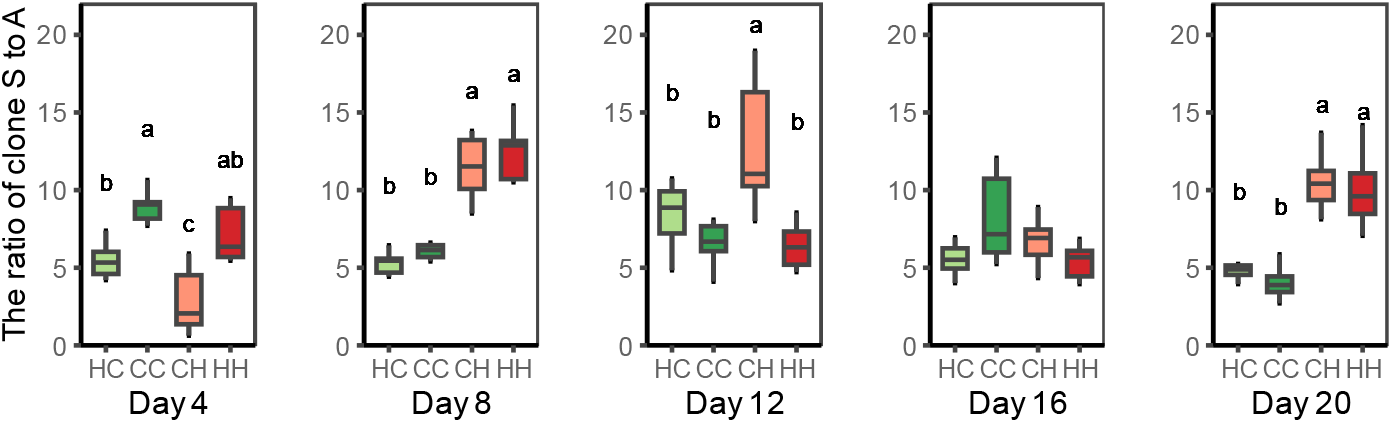
Proportions of S/A clones determined by COI metabarcoding across five temporal sampling points under maternal high temperature exposure. The two-letter codes in X-axis labels: the first capital letter indicates the maternal environment, while the second indicates the offspring environment. C represents control temperature (20°C), and H represents high temperature (28°C).

### 3.3 Impact of clonal maternal low-temperature exposure on mixed-clone population dynamics and genetic structure

In low-temperature exposure experiment, statistical analysis indicated that offspring environmental temperature significantly influenced mixed clonal population density (P < 0.001,Table 3), mictic ratio (P = 0.032,Table 3), and daily resting eggs production (P <0.001,Table 3). Maternal environmental temperature showed no significant effect on these parameters, and there was no interaction between the maternal and offspring environmental temperatures (Table 3). Groups with identical offspring temperatures (CC and LC, LL and CL) displayed greater similarity in the temporal trends of mixed clonal population density, mictic ratio, and daily resting eggs production (Fig. 5). Groups CC and LC reached their peak population densities on days 17 and 19, respectively, followed by a decline, and consistently maintained higher densities than groups CL and LL, which were slowly increasing throughout the experimental period (Fig. 5a). Mictic ratio in the CC and LC groups peaked early (within the first 6 days) followed by a decrease (Fig. 5b).

**Table 3.** Results of the GEE two-way ANOVA on quantitative dynamic characteristics (population density, mictic ratio, daily resting eggs production) of mixed clonal populations in the lox-temperature maternal effects experiment.

|  | a) Population density |  |  |  | b) Mictic ratio |  |  |  | c) Number of resting eggs |  |  |  |
| --- | --- | --- | --- | --- | --- | --- | --- | --- | --- | --- | --- | --- |
|  | Estimate | Std Error | Z value | P value | Estimate | Std Error | Z value | P value | Estimate | Std Error | Z value | P value |
| Maternal environment(A) | -1.830 | 4.700 | 0.152 | 0.697 | 0.005 | 0.027 | 0.039 | 0.844 | 4.430 | 9.570 | 0.214 | 0.644 |
| Offspring environment(B) | -31.530 | 3.520 | 80.251 | <0.001 | -0.044 | 0.021 | 4.577 | 0.032 | -31.750 | 5.640 | 31.631 | <0.001 |
| A×B | 1.930 | 4.710 | 0.167 | 0.683 | 0.018 | 0.032 | 0.311 | 0.577 | -4.350 | 9.570 | 0.206 | 0.650 |

**Fig. 5.**
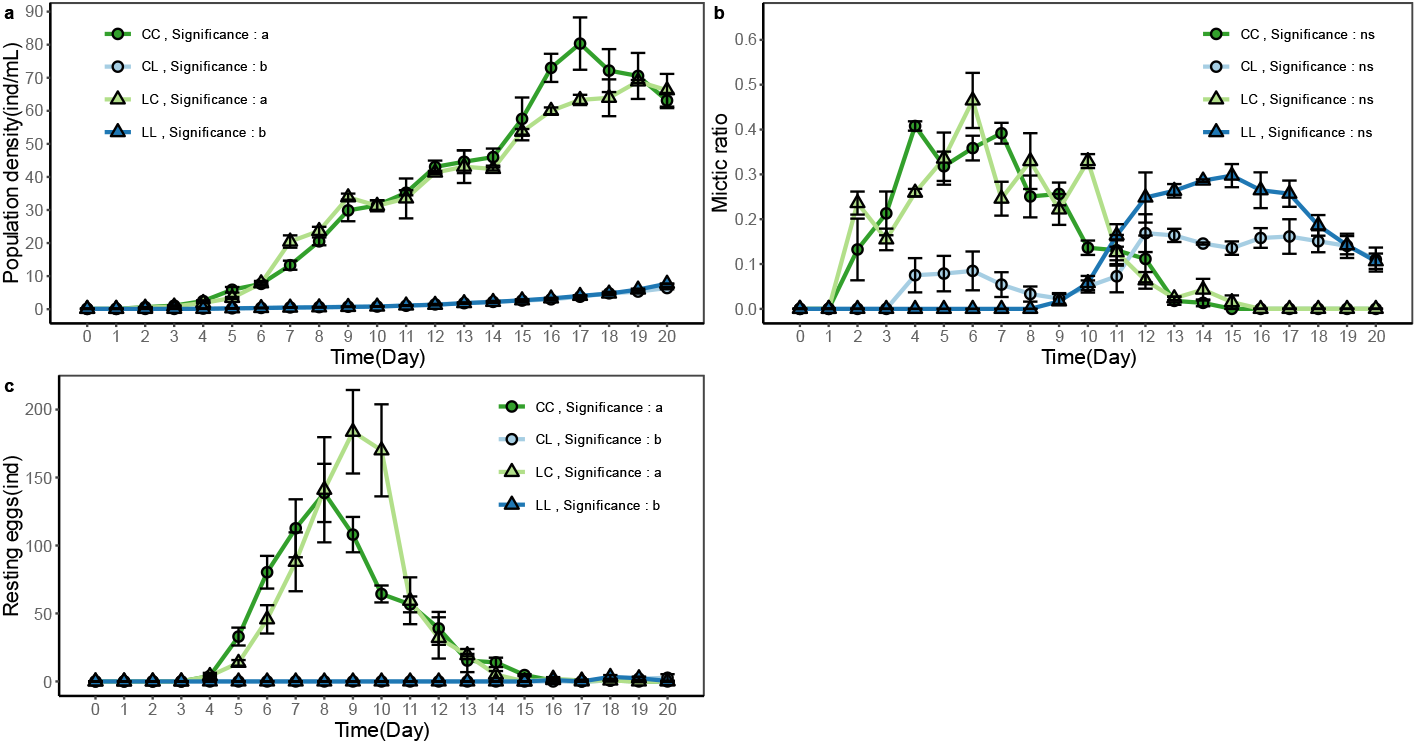
Population dynamics of mixed-clone populations under maternal cold effects in low-temperature experiments. a: Temporal changes in population density; b: Mictic ratio (= mictic females / (mictic females + amictic females)); c: Daily resting eggs production. In the two-letter codes, the first capital letter indicates the maternal environment, while the second indicates the offspring environment. C represents control temperature (20°C), and L represents low temperature (12°C). Curves labeled with different lowercase letters differ significantly (P < 0.05) as determined by GEE modeling. Data presented as mean ± SE.

The CL group showed an initial small peak and decline in mictic ratio within the first 6 days, subsequently forming a higher second peak (Fig. 5b). Mictic females were initially absent in the early of LL group, and the mictic ratio in LL commenced rising coinciding with the second peak in CL (Fig. 5b). For resting eggs production, the CC and LC groups peaked in day 8 and day 9, respectively, followed by a reduction. Groups CL and LL produced almost no resting eggs during the experimental period (Fig. 5c).

Results of the maternal cold experiment showed that clonal ratios were not affected by maternal environmental temperature at any of the five time points (P > 0.05,Table 4), whereas they were significantly influenced by the offspring’s environmental temperature (P < 0.001,Table 4), moreover, there were clear interactions between the two factors on days 4, 8, and 16 (P < 0.05,Table 4). Clonal ratio in the LL group was significantly higher than that in the CL group on day 4, then the difference gradually diminished on days 8 and 12 to none. Conversely, on days 16 and 20, the CL group showed a significantly higher clonal proportion than the LL group (Fig. 6). Clonal ratio in the CC group was significantly higher than that in the LC group on days 4 and 8, but on day 16, the clonal ratio in LC group became clearly higher than that in CC group. By day 20, no significant difference was found between the CC and LC groups (Fig. 6).

**Table 4.** Results of two-way ANOVA on the proportional representation of S-clone and A-clone COI metabarcoding sequences at five time points in the low-temperature maternal effects experiment. F-value represents the parametric two-way ANOVA statistic, whereas H indicates the non-parametric Scheirer-Ray-Hare.

|  | a) Ratio of clone S to A in day 4 |  |  |  | b) Ratio of clone S to A in day 8 |  |  |  | c) Ratio of clone S to A in day 12 |  |  |  |
| --- | --- | --- | --- | --- | --- | --- | --- | --- | --- | --- | --- | --- |
|  | df | Sum sq | F value | P value | df | Sum Sq | H | P value | df | Sum Sq | H | P value |
| Maternal environment(A) | 1 | 0.103 | 1.576 | 0.218 | 1 | 44.444 | 0.400 | 0.527 | 1 | 87.111 | 0.785 | 0.376 |
| Offspring environment(B) | 1 | 0.791 | 12.127 | 0.001 | 1 | 2401.000 | 21.631 | <0.001 | 1 | 2916.000 | 26.270 | <0.001 |
| A×B | 1 | 1.863 | 28.549 | <0.001 | 1 | 641.778 | 5.782 | 0.016 | 1 | 49.000 | 0.441 | 0.506 |
| Residuals | 32 | 2.088 |  |  | 32 | 797.778 |  |  | 32 | 832.889 |  |  |
|  | d) Ratio of clone S to A in day 16 |  |  |  | e) Ratio of clone S to A in day 20 |  |  |  |  |  |  |  |
|  | df | Sum Sq | H | P value | df | Sum Sq | H | P value |  |  |  |  |
| Maternal environment(A) | 1 | 0.000 | 0.000 | 1.000 | 1 | 245.444 | 2.211 | 0.137 |  |  |  |  |
| Offspring environment(B) | 1 | 2240.444 | 20.184 | <0.001 | 1 | 2916 | 26.27 | <0.001 |  |  |  |  |
| A×B | 1 | 1133.444 | 10.211 | 0.001 | 1 | 128.444 | 1.157 | 0.282 |  |  |  |  |
| Residuals | 32 | 511.111 |  |  | 32 | 595.111 |  |  |  |  |  |  |

**Fig. 6.**
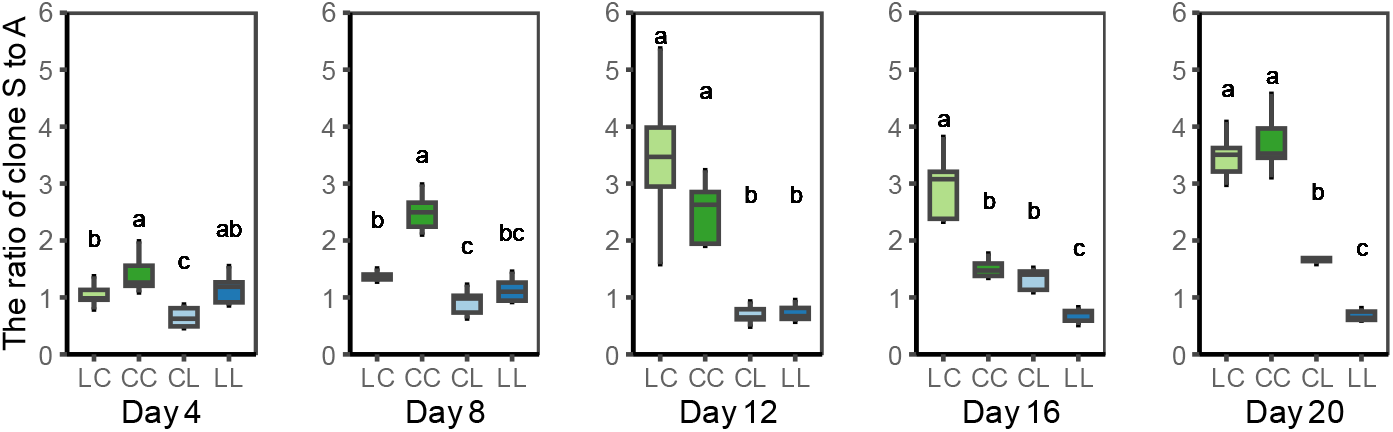
Proportions of S/A clones determined by COI metabarcoding across five temporal sampling points under maternal cold exposure. The two-letter codes in X-axis labels: the first capital letter indicates the maternal environment, while the second indicates the offspring environment. C represents control temperature (20°C), and L represents low temperature (12°C).

## 4 .Discussion

In this study, we observed that the relative proportion of S clones experiencing matched maternal and offspring temperatures (i.e., S clones in LL, CC, HH groups) was significantly greater than those experiencing mismatched maternal-offspring temperatures (i.e., S clones in CL, LC, CH groups) during the initial period (Day 4: LL > CL, P < 0.05; HH > CH, P < 0.05; Days 4 & 8: CC > LC, P < 0.05; Fig. 4, Fig. 6).Current evidences across diverse species have demonstrated that non-lethal heat exposure in maternal generation can significantly improves offspring performance under high-temperature (Garbutt *et al*., 2014, Salinas & Munch, 2012, Sun *et al*., 2023), and parental cold exposure improves offspring low-temperature fitness(Shama *et al*., 2014, Stelzer, 2002, Sun & Niu, 2012). For example, in marine sticklebacks, mothers modulate offspring mitochondrial capacity and efficiency to confers a growth advantage in body size through “metabolic matching” in thermal environments (Shama *et al*., 2014). Consistent with these findings, our results clearly indicate that the matching of maternal and offspring environments, mediated by transgenerational effects, significantly enhanced the fitness of clone S during early competition, thereby rapidly altering the population genetic composition towards a bias favoring clone S.

More interestingly, we found that the competitive advantage conferred by maternal effects has a distinct temporal pattern. Notably, the early proportional advantage of clone S was systematically reversed in several treatment combinations: Under high temperature, clone S’s advantage in the HH group at day 4 was surpassed by the CH group by day 12 (P < 0.05), and subsequently, differences between groups disappeared (Fig. 4;Fig. 7). Under low temperature, the initial advantage held by the LL group over CL at day 4 (P < 0.05) was reversed during days 16-20 (CL group exceeded LL, P < 0.05,Fig. 6;Fig. 7). While, in the control temperature treatments (CC vs. HC, CC vs. LC), the early advantage of the CC group (CC > HC on day 4,Fig. 4; CC > LC on days 4 and 8,Fig. 6; P<0.05;Fig. 7) diminished in later stages. This reversal in clonal proportion strongly indicates that changes in genetic structure induced by maternal effects are dynamic and non-persistent.

**Fig. 7.**
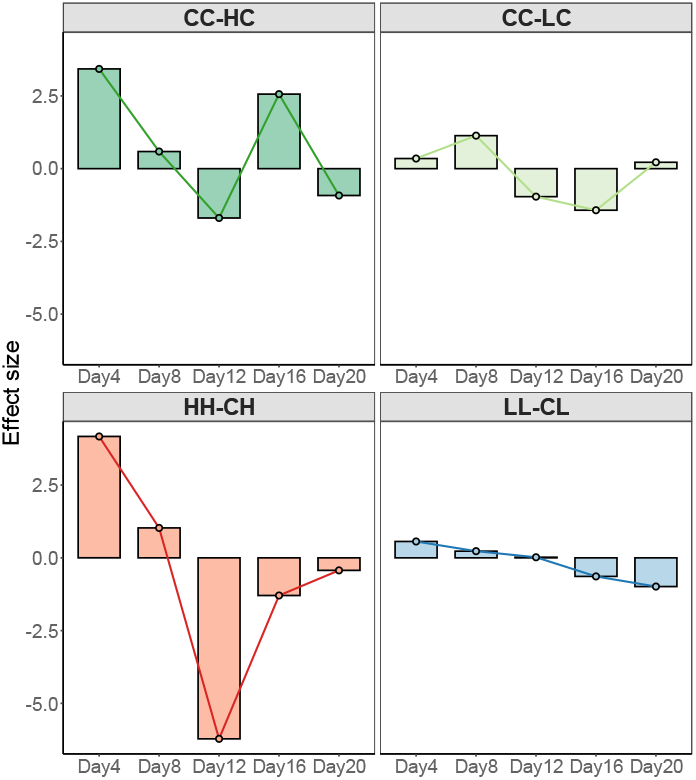
The effect size of maternal effects on clone proportions. Bar heights and data point values indicate the difference in clone proportions between the maternal effect treatment groups and the non-maternal effect control groups (e.g., CC-HC, CC-LC, HH-CH, LL-CL), where positive values indicate that the maternal effect confers a competitive advantage, and negative values signify the attenuation or loss of this advantage.

Potential mechanisms underlying this temporal limitation and reversal may be: (1) The decay of epigenetic modifications during transmission across generations(Feiner *et al*., 2022, Khan & Mishra, 2024, Tariel *et al*., 2020), leading to gradual loss of maternally pre-programmed information; (2) The accumulation of adaptive costs incurred across multiple generations under sustained stress, which counteracts the initial advantages(Carvalho *et al*., 2023, Tariel *et al*., 2020). Integrating these broader findings, the reversal in clonal ratio dominance observed in our experiment is highly likely the result of the gradual dissipation or even reversal of the adaptive maternal temperature effect across generational transmission. With the diminishment of maternal effect, offspring phenotype and fitness are progressively dominated by the conditions of their own immediate environment, ultimately leading to a convergence in clonal ratio among different maternal treatment groups when cultured under the same environment.

Environmental temperature not merely act as a selective pressure, but also could profoundly influence the temporal trajectory of these clone competitive dynamics and consequently population genetic structure alterations, through its regulation of population metabolism and generational succession rates. The low-temperature experimental groups (LL vs CL) and control-temperature groups (CC vs LC) displayed the same “maternal-effect advantage → advantage attenuation/reversal → environmental dominance” pattern as the high-temperature group (HH vs CH), though their dynamic processes were temporally extended: The advantage in the LL group persisted until days 16-20 before reversal occurred, whereas the advantage of the CC group under control temperature disappeared after day 8 (with no significant reversal observed, Fig. 6;Fig. 7). Reason for this difference lies in temperature’s strong modulation of rotifer life-history characteristics—higher temperatures substantially elevate metabolic rates, resulting in faster reproduction, reduced longevity, and accelerated generational succession (Gillooly, 2000, Sun & Niu, 2012, Walczyńska *et al*., 2017).

Thus, our results show that temperature serves not only as an environmental selective pressure but also determines the temporal scale over which maternal effects influence competitive dynamics. Under high temperature with vigorous metabolism, alterations in genetic structure induced by maternal effects occur rapidly and dissipate quickly, while under low temperature, this transformation process is markedly prolonged. This mechanistic framework explains the timing differences in clone S/A ratio dominance and reversal events across temperature treatments. While significant proportion reversal was absent in the low-temperature experiment’s control temperature group (CC vs LC), a trend of higher LC proportion than CC presented on day 12 (despite p>0.05,Fig. 6). Meanwhile, in the high-temperature experiment’s control group (CC vs HC), maternal effect advantage was only apparent at day 4,with no statistically significant differences detected subsequently (Fig. 4), but the directional patterns conform to our proposed pattern. These findings imply possibly missed peaks because of restricted sampling frequency, providing additional support for the metabolic rate-driven maternal effect temporal dynamic model. A higher frequency of high-throughput sequencing can be employed in future research to delineate the regulatory mode of maternal effects on population genetic structure with greater precision.

In addition, precise quantification of clonal competition dynamics in mixed populations is a fundamental prerequisite for understanding how maternal effects reconfigure population genetic structure. This study successfully established a high-throughput monitoring system based on COI metabarcoding. Systematic validation using artificially mixed populations of known proportions confirmed the exceptionally high reliability of this method for quantifying rotifer clonal proportions (R^2^ > 0.975, P < 0.001), with stable accuracy maintained across a 97%-100% similarity threshold range (Fig. 2). It should be noted that the high quantitative accuracy achieved here was benefited from the substantial COI genetic distance (>0.05) between clones S and A, and the custom reference database constructed from these two clones, which effectively reduced interference from sequence homology and background noise. The primers targeted regions with high single-nucleotide polymorphism (SNP) divergence and have been previously validated as effective universal primers for zooplankton(Elbrecht & Leese, 2017, Leray *et al*., 2013), ensuring high amplification specificity. Consequently, three critical prerequisites ensure the reliability of DNA metabarcoding for quantifying clones at the sub-species level: adequate inter-clonal genetic distance, high-performance primer amplification, and a population-specific exclusive reference database. This provides a clear optimization pathway for future extension of this technology to more complex multi-clonal systems.

In conclusion, this study established a clonal dynamic monitoring system of monogonant rotifer based on COI metabarcoding, revealing that maternal temperature effects can significantly alter the early competitive fitness of clones, leading to short-term restructuring of population genetic structure. Yet, with the intergenerational decay of maternal effects, this clonal superiority undergoes reversal and gradual dissipation, whereas selection on inherent fitness mediated by ambient temperature increasingly governs the stability of competitive outcomes. A clear temporal threshold characterizes this effect: elevated temperatures shorten its duration, accelerating the return to environmentally-determined genetic structure, while lower temperatures prolong its persistence. This study provides both a methodological foundation and empirical support for high-resolution monitoring of rotifer population dynamics at the intraspecific level, holding significant implications for understanding population genetic dynamics under maternal effects.

